# Anomalous diffusion in inverted variable-lengthscale fluorescence correlation spectroscopy

**DOI:** 10.1101/359240

**Authors:** M. Stolle, C. Fradin

## Abstract

Using fluorescence correlation spectroscopy (FCS) to distinguish between different types of diffusion processes is often a perilous undertaking, as the analysis of the resulting autocorrelation data is model-dependant. Two recently introduced strategies, however, can help move towards a model-independent interpretation of FCS experiments: 1) the obtention of correlation data at different length-scales and 2) its inversion to retrieve the mean-squared displacement associated with the process under study. We use computer simulations to examine the signature of several biologically relevant diffusion processes (simple diffusion, continuous-time random walk, caged diffusion, obstructed diffusion, two-state diffusion and diffusing diffusivity) in variable-lengthscale FCS. We show that, when used in concert, lengthscale variation and data inversion permit to identify non-Gaussian processes and, regardless of Gaussianity, to retrieve their mean-squared displacement over several orders of magnitude in time. This makes unbiased discrimination between different classes of diffusion models possible.

## 1 Introduction

Quantifying the motions of macromolecules in cells, while an important task, is complicated. The cellular environment is crowded and heterogenous, and many biomolecules transiently interact with others. Because of this, macromolecular diffusion in cells can take many forms and is seldom simply Brownian. It may exhibit a mean-squared displacement (MSD) that is not linear in time, a distribution of displacements that is not Gaussian, or both (1). Langowski’s study of the diffusion of the green fluorescent protein in cells, using fluorescence correlation spectroscopy (FCS), was one of the first to put this anomalous behaviour in the spotlight (2).

FCS, a technique based on the statistical analysis of the fluorescence signal recorded from a well-defined observation volume (radius*w)* is perfectly suited to the range of concentrations (1 to 100 nM) and diffusion coefficients (1 to 100 *µ*m^2^/s) typically encountered for soluble proteins in cells (3, 4). Further, the systematic variation of w has made this technique especially useful for the study of anomalous diffusion processes, which are often lengthscale-dependent. This scheme, known as spot variation FCS or more generally variable-lengthscale FCS (VLS-FCS), allows the construction of the so-called “diffusion law”, that is the relationship between projected area of the observation volume, *w*^2^, and characteristic decay time of the FCS autocorrelation function (ACF), *Τ*_1/2_ (5). The diffusion law provides a proxy for the particles MSD. Initially, VLS-FCS was achieved by enlarging the usually diffraction-limited point-spread function (PSF) that sets the value of w (5–7). Since then, several methods have been proposed to extend the range of available w below the diffraction limit (8–10). Another interesting development has been the generation of VLS-FCS data from imaging modalities (total internal reflection, single-plane illumination), in which case the observation volume size can be varied by binning pixels (11, 12).

The ultimate limit on FCS spatial resolution, however, is not set by the size of the observation volume, but by the temporal resolution of the signal from which the ACF is computed (13–15). The displacements of fluorophores over distance smaller than *w* still result in changes in fluorescence intensity - with low contrast -, and they are therefore captured in the short-time regime of the ACF. Thus even with a fixed-size observation volume, FCS can resolve motions over a range of lengthscales on either sides of the diffraction limit. This capacity was demonstrated in a cluster of studies exploring DNA segments dynamics, where the ACF was inverted to directly recover the MSD of the segments (16–18). The same ACF inversion procedure was later used to study the effect of crowding in membranes and polymer solutions (19, 20). A similar idea was later implemented for image correlation spectroscopy, using a different mathematical scheme to extract the MSD from spatiotemporal ACFs, and used to characterize the anomalous behaviour of GFP diffusion in cells (15, 21). Of note, both these MSD recovery schemes work under the assumption of a process with a Gaussian propagator.

Following analytical work by Höfling & Franosch (1, 22), we recently showed that variation of the FCS observation volume size can be combined with ACF inversion to obtain the MSD of tracer particles for over 5 orders of magnitude in time (20). The two model systems we studied (crowded dextran solutions and agarose gels) behaved very differently in regard to the superimposition of the apparent MSDs extracted from ACFs obtained at different length-scales (20). This dissimilarity reflects a qualitative difference in the nature of the propagator (a.k.a. distribution of displacements) of the underlying process. Since only processes with Gaussian propagators are expected to lead to a perfect superimposition of the apparent MSDs, the combination of VLS-FCS with ACF inversion provides a test of the Gaussianity of the diffusive process (20, 23).

Here we study the signature of different biologically relevant diffusive processes (continuous time random walk, two-component diffusion, diffusing diffusivity, obstructed diffusion, caged diffusion). Since analytical solutions are available only in a small number of limiting cases, we performed 3D single agent simulations for all these processes, and obtained apparent MSDs from the inversion of ACFs simulated for a range of *w*. By highlighting the different signatures of each diffusion process, our work provides a benchmark for model-independent interpretation of inverted VLS-FCS experiments. It also refines our understanding of the relationship between apparent MSD and real MSD. We show in particular that the inversion procedure yields the correct MSD at short lag times regardless of the nature of the propagator, suggesting possible refinements for VLS-FCS experiments.

## 2 Theory

### 2.1 General form of the ACF

The fluorescence signal *I(t)* obtained in an FCS experiment is correlated in time to give the ACF, 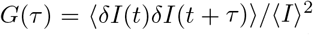, where 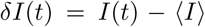. For a transport process with propagator 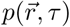 and a 3D Gaussian observation volume (radius *w*, aspect ratio *S*) with normalized profile 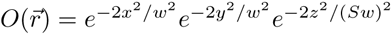, the ACF becomes:

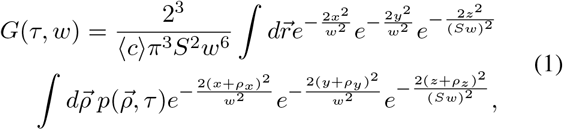

where (*c*) is the average fluorophore concentration.

### 2.2 ACF for an isotropic Gaussian diffusive process

If the propagator is both Gaussian and isotropic, it can be expressed as a function of the MSD, {*r*^2^} (*τ*):

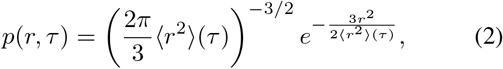

which leads to:

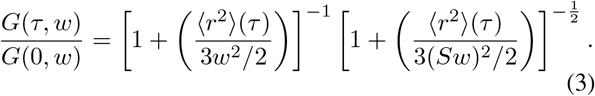

The above equation is valid for any lag time T at which the propagator is Gaussian. It is the basis for the inversion procedure used in this work, and illustrated in Fig. 1, where 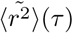 is obtained from *G(Τ, W)/G*(0, *w*) by inverting Eq. 3 (the tilde is used as a reminder that the apparent MSD extracted from the ACF might differ from the actual MSD when the propagator is not Gaussian).

**Figure 1:**
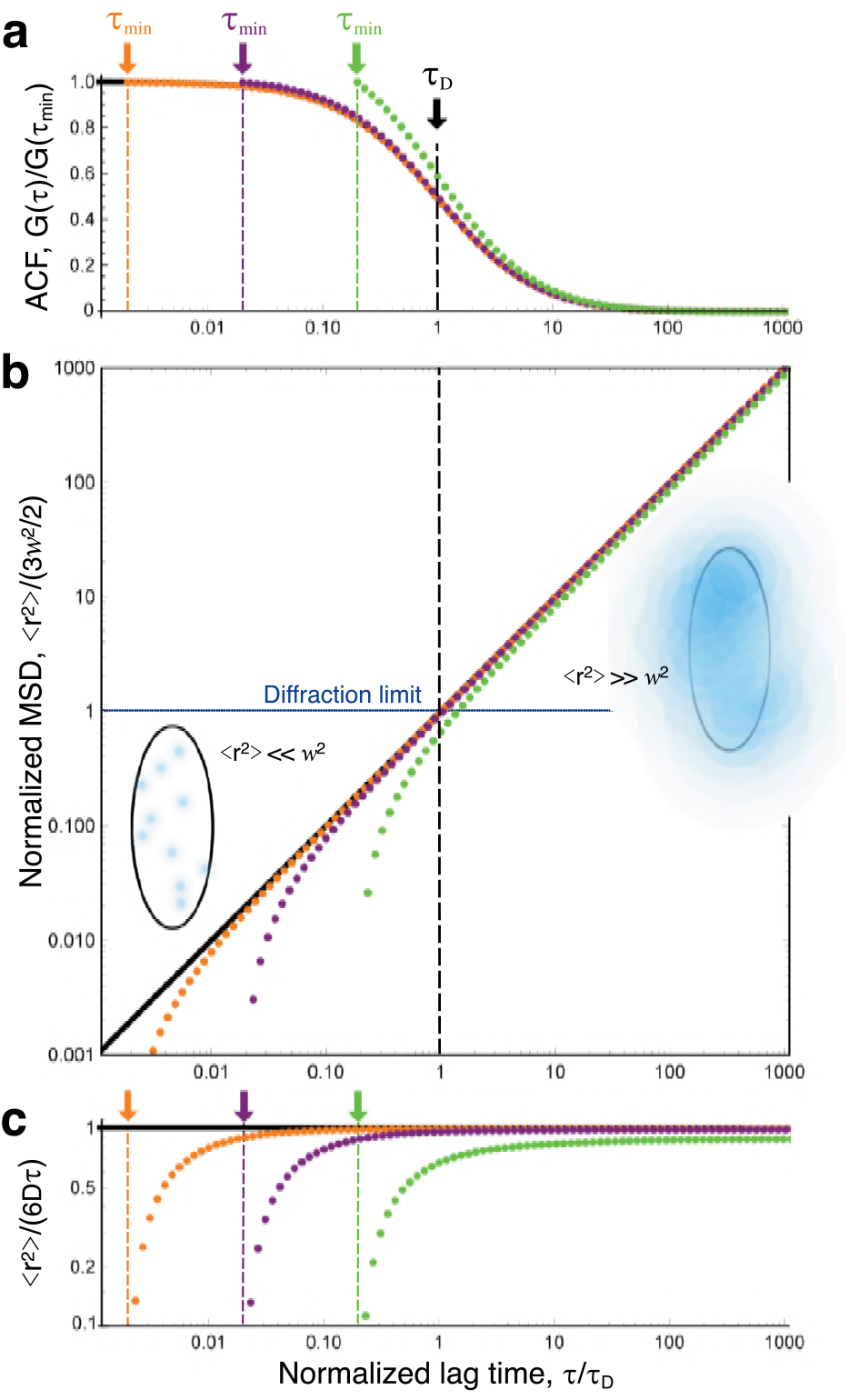
(a) Simple diffusion ACFs (analytical form), normalized by amplitude at shortest lag time (black line: *τ*_min_ = 0, orange symbols: *τ*_min_ = *τD*/500, purple symbols: *τ*_min_ = τD/50, green symbols: *τ*_min_ = *τ*D/50). (b) Apparent MSDs obtained by inversion of the ACFs shown in (a), showing a departure from the real MSD at lag times close to *τ*_min_. Schematic representations of the detection volume illustrate particle displacements shorter and larger than the diffraction limit, for lag times below and above *τ_D_*, respectively. (c) Apparent and actual MSDs after normalization by 6 *Dτ*.

### 2.3 Relationship between ACF and MSD at short lag times

We show in this section that for short *τ* there is a simple linear relationship between *G(Τ, W)/G*(0, *w*) – 1 and (*r*^2^)(*r*), whether the propagator is Gaussian or not. The only necessary assumption is that the propagator is isotropic, in which case it can be written *p(r, τ)*. At lag times much smaller than the ACF characteristic decay time 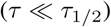, particles have not yet diffused over distances comparable to the observation volume radius, in other words 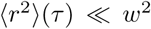. This means that*p(r*, *τ*) ~ 0 for *r* > *w*, in which case 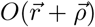 can be replaced in Eq. 1 by the first two even terms of its Taylor series expansion in *ρ/w* (the odd terms are left out as they disappear when integrating over 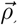). Performing the integration over 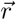 in Eq. 1 then yields:

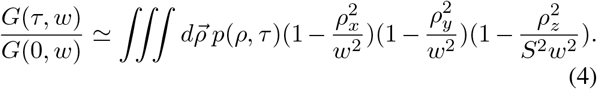

Again ignoring higher order terms in (*ρ*/*w*)*^2^*, and using the 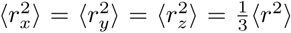 valid for an isotropic propagator, we get (for 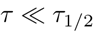, or 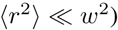:

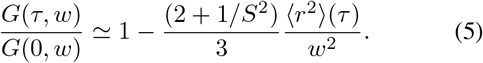

For a Gaussian process, this linear relationship between ACF and MSD can be recovered directly from Eq. 3, by performing a first-order Taylor expansion in (*r*^2^)/*w*^2^. A more general form of this equation has been derived in Ref. (24).

### 2.4 Normalization of the ACF

When inverting an experimentally obtained ACF to obtain the apparent MSD, the first step is to normalize its amplitude to obtain *G(Τ, W*)/*G* (0, *w*). When the actual value of *G* (0, *w)* is unknown, the most straightforward solution is to perform the inversion on *G(τ, w*)/*G*(*τ*_min_, *w*), where *τ*_min_ is the shortest lag time at which a reliable value of the ACF can be obtained (Fig. 1). Obviously, for the inversion to work properly, one needs 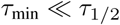. If this condition is fulfilled, then at short *τ* (when Eq. 5 is valid) the apparent MSD is:

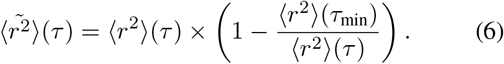

This highlights an additional necessary condition for the inversion procedure to work properly, which is that *τ* needs to be large enough for 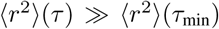. This is illustrated in Fig. 1 for a simple diffusive process: 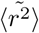 deviates from the actual 〈*r*^2^〉 by less than 5% as long as *τ* > 20 *τ* _min_.

### 2.5 ACF for a truncated detection volume

In the simulations presented below, the observation volume profile was truncated in all three space directions to reduce computational times. Truncated observation volumes have been considered before, for either reflective or absorbing boundary conditions (25, 26). Here we consider the case that corresponds to our simulations, where particles become invisible when leaving the rectangular volume with dimensions 2 *bw* x 2 *bw* x 2 *bSw* centred on the observation volume, but allowed to diffuse in and out. For a Gaussian propagator and Gaussian detection profile, the ACF is then given by:

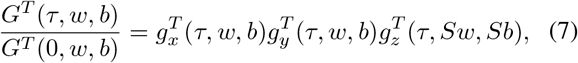

where:

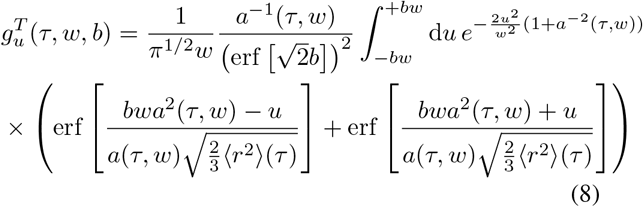

and:

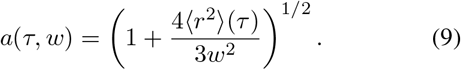

Although there is no simple analytical expression for *G^T^ (τ, w, b)*, Eq. 8 can be integrated numerically. The truncated ACF (Eq. 7) becomes indistinguishable from the ACF obtained in the absence of truncation (Eq. 1) for *b* ≥ 1 (Fig. 2). Since in our simulations *b* = 7.5, we consider in the following that *G^T^(τ, w, b) ~G*(*τ*, *w*).

**Figure 2:**
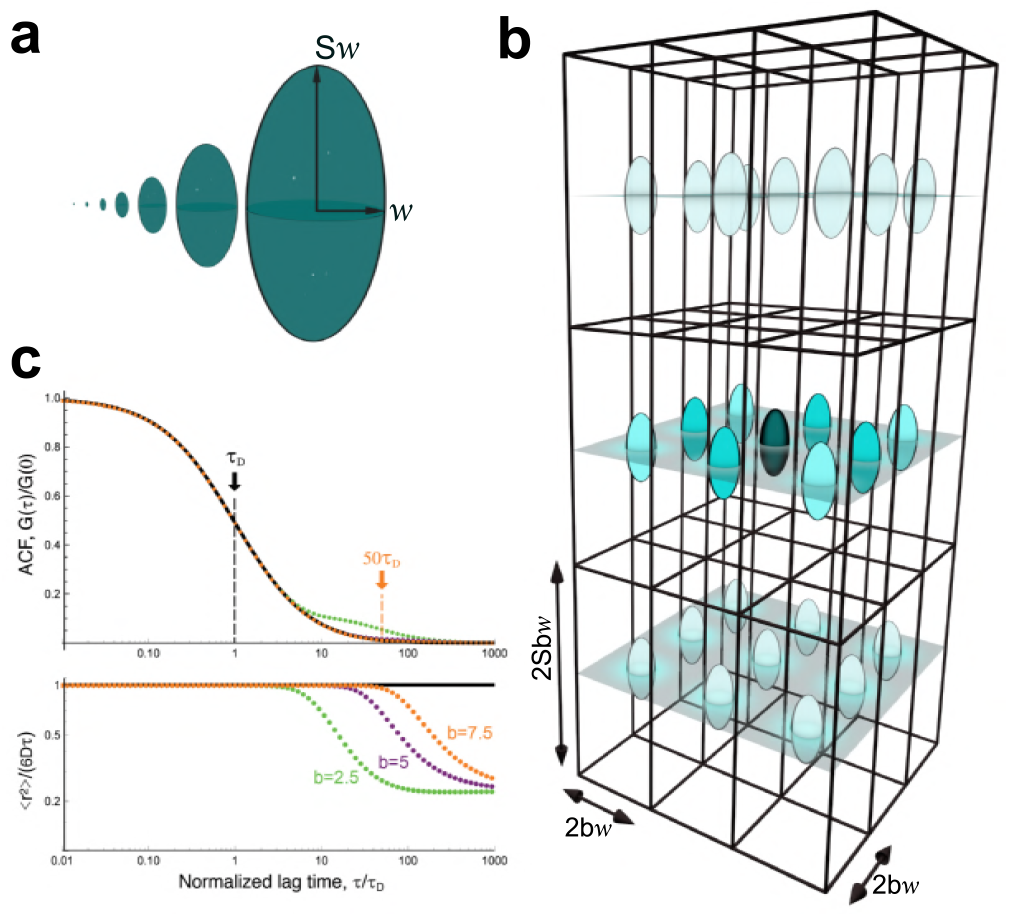
(a) Respective sizes of the 7 different 3D-Gaussian ellipsoidal observation volumes used in our simulations. (b) Array of observation volumes. (c) ACFs expected for simple diffusion in a single observation volume (black line, Eq. 3) or for a regular array of detection volumes (green symbols:*b =* 2.5, purple symbols: *b* = 5, orange symbols: *b* = 7.5, Eq. 11). The lower panel shows the corresponding MSD *((r*^2^)/(6*Dτ*) as obtained by inversion of the ACFs shown in the upper panel).

### 2.6 ACF for a regular array of detection volumes

To make our VLS-FCS simulations more efficient, we calculated the signal from an array of observation volumes spaced by*2bw* in the focal plane and *2bSw* along the optical axis (Fig. 2). In this case the ACF takes the form:

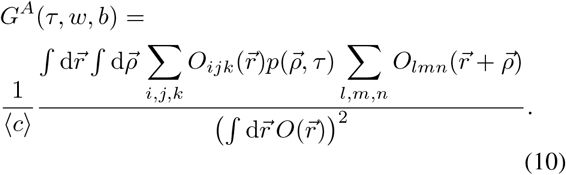

where 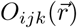 is the intensity profile of the detection volume centred at {2 *bwi*,*2bwj, 2bSwk}.* Eq. 10 can be simplified by recognizing that only adjacent detection volumes are likely to record correlated events. Thus for a *n* × *n* × *n* array:

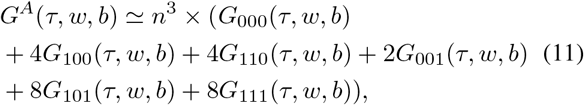

where *G_opq_* denotes the correlation between two detection volumes spaced by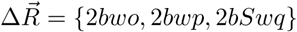:

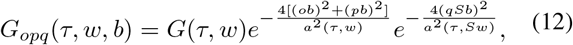

where *a^2^(τ,w)* is given by Eq. 9. This expression can be derived in the same way as the ACF for two-focus FCS experiments (27). For an elongated observation volume (*S* = 5), the first three terms in Eq. 11 dominate, where the maxima for *G*_100_ (*τ*, *w, b)* and *G*_110_(*τ*, *w, b)* are much higher and occur at shorter lag times than for the other terms. As can be expected, 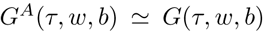 for 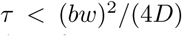 (Fig. 2c). Thus in practice simulations done for an array of detection volumes can be used for ACF inversion up to 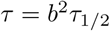 (in our case, 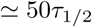).

## 3 Methods

### 3.1 Generation of particle trajectories

We used a generic 3D single-agent continuum simulation procedure, written in C++, to generate particle trajectories and simulate VLS-FCS experiments with observation volume varying in size between*w* = 0.3 and 30 *µ*m. Particles (usually 32) were placed at random positions within a 3D rectangular simulation box with periodic boundary conditions. The box dimensions,*a x a x* 5 a with*a* = 450 *µ*m, were chosen to be 15 times larger than those of the largest observation volume considered. A time step Δ*t* = 100 ns was selected to ensure particle trajectories with sufficient resolution, even in the smallest observation volume considered. Simulations were typically run for 10 ^11^ time steps. For simple diffusion, displacements at each step and in each spatial directi<u>on wer</u>e drawn from a normal distribution with variance 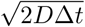 (diffusion coefficient*D* = 500 *µ*m^2^/s).

### 3.2 Anomalous and obstructed trajectories

The algorithm used to simulate simple Brownian diffusion was modified to simulate 5 different diffusive processes, illustrated in Fig. 3.*i)* Simulations of continuous random walks (CTRW) were carried out following Refs. (28–30). Step lengths were drawn from the same Gaussian distribution as for simple diffusion, but after each step a wait time τ_W_ was added, with duration drawn from a Pareto distribution 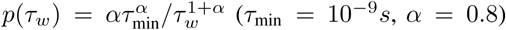. To allow for a range of wait times, the time step was changed for this diffusion process to 10^−9^*s* and simulations run for 10 ^13^ steps.*ii)* Two-component diffusion was simulated by allowing tracer particles to switch between a fast*(D* = 500 *µ*m^2^/s) and a slow diffusive state (D*’* = 50 *µ*m^2^/*s*). Transitions between states were assumed to be Poisson processes with constant rates, *k_on_* = *k_off_* = 0 or *k_on_* = *k_off_* = 500 s^−1^.*iii)* Diffusing diffusivity was simulated following Chubynsky & Slater (31). The diffusion coefficient of each particle was allowed to undergo a 1D random walk between *D*_min_ = 0 *µ*m^2^/s and *D*_max_ = 500 *µ*m^2^/s (with a random initial value within that range), with a “diffusion coefficient” *d* = 5.5 × 10 ^−17^ *m*^4^/*s*^3^, chosen such that the full range of possible diffusion coefficients was explored by a given particle over a time comparable to its characteristic diffusion time through one of the smaller observation volumes.*iv)* The obstructed diffusion of particles hindered by the presence of fixed obstacles was simulated as previously done in 2D by Saxton (28). Fixed reflective cubic obstacles with dimensions 0.75 *µ*m were placed randomly on a cubic lattice, at a volume fraction *$* = 0.310 just below the percolation limit (* = 0.3116). *v)* Caged diffusion in cubic 1 *µ*m corrals separated by semi-permeable barriers was simulated as done in 2D in Refs. (5, 32). Particles could cross barriers only with probability *p* = 0.005 and were reflected otherwise. For the obstructed and caged diffusion models, the value of *b* was changed very slightly for the smallest observation volumes, to avoid always placing their center in the same position within the cubic cells used to generate the obstacles.

**Figure 3:**
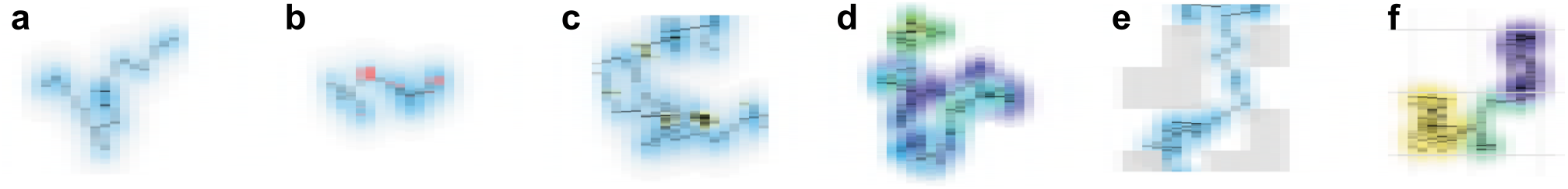
Illustration of the different models considered, showing short two-dimensional trajectories. Successive positions are marked by black dots, while pink dots signal a wait time before taking the next step. The halos around each position represent the Gaussian distribution from which the next step is drawn. The colour of the halos represent either the value of the diffusion coefficient (a-e) or in which cage the particle is at that time (f).

### 3.3 MSD and non-Gaussian parameter

Both the second moment ((*r*^2^), mean-squared displacement) and fourth moment ((*r*^4^)) of the particle displacement were calculated from particle trajectories using two separate sampling times. For the first 10 ^7^ steps, particle positions were saved at each step (every 100 ns), and from these (*r*^2^)(*τ*) and (*r*^4^) (*τ*) calculated up to *τ* = 1 *s*. After 10^7^ steps, positions were saved only every 1 ms, and (*r*^2^)(*τ*) and (*r*^4^)(*τ*) were calculated from this parsed data for *τ* > 1 ms. The moments calculated with different sampling times were stitched together to give data sets spanning the full explored time range, *τ* = 100 ns to 10 ^4^ s. The non-Gaussian parameter, *β*, was then calculated according to its definition:

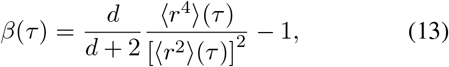

where *d* = 3 is the number of spatial dimensions (1).

### 3.4 Computation of the ACFs

A set of 7 ACFs was calculated from the same particle trajectories, for 3D Gaussian detection volumes with 1 /*e*^2^ radii, *w_i_* (*i* =1 to 7), ranging from 300 nm to 30 *µ*m and equally spaced on a log scale. The aspect ratio was kept the same for all detection volumes, *S* = 5. In order to increase the number of particle trajectories passing through the smaller detection volumes, the simulation box was split into smaller boxes of size 2 *bw_i_* x 2 *bw_i_* x 2 *bSw_i_*, with *b* = 7.5, and a detection volume placed at the centre of each of these smaller boxes (Fig. 2b). For each *w_i_*, the fluorescence intensity collected for a particle at (*x*,*y*,*z*) was thus calculated as:

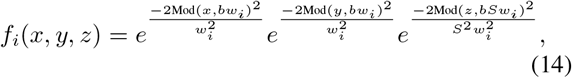

where Mod(*x*, *bw_i_*) is the remainder of the division of *x* by *bw_i_*. At each simulation step, *s*, and for each detection volume size, *w*_i_, the fluorescence intensity emitted by each particle was calculated using Eq. 14, and individual particle intensities were summed to give the total intensity, *F_i_* (*s ΔΤ*). Up to 10 ^7^ steps (1 s), the simulated intensity was saved at each step, after which only binned data (1 ms bins) was saved. At the end of the simulation, both the original and binned data were correlated using a discrete fast Fourier transform, to obtain the ACF between 10^−7^ and 1 s, and 1 ms and 10^3^ s, respectively. A symmetric normalization procedure was used to correct for the discrete nature of the correlation, imposing that the numerator and denominator of the ACF are calculated from the same number of points (33).

### 3.5 ACF inversion

An apparent MSD, 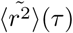, was calculated from each ACF using an inversion procedure justified in the case of Gaussian diffusion (16, 19). The amplitude of the ACFs was first normalized to 1 using *G*(*ΔΤ*, *w*), after which Eq. 3 was solved for (*r*^2^) at each lag time, *τ*, to obtain 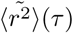.

ACFs were also fitted to the general expression derived for a Gaussian anomalous process with 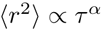 (1, 34):

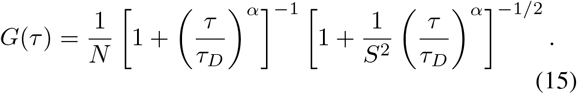

## 4 Results

We performed 3D single-agent simulations for different diffusive processes and characterized the obtained particle trajectories in two different ways. First, we calculated the mean-squared displacement of the particles, (*r*^2^)(*τ*), the non-Gaussian parameter, *β*(*τ*), and the distribution of displacements, *P*(*x*, *τ*), directly from the trajectories (represented by black lines or symbols in Figs. 4–12). Second, we simulated the result of VLS-FCS experiments, generating ACFs for observation volumes ranging in size from *w* = 0.3 to 30*µ*m (ACFs and derived parameters are represented by colored symbols in Figs. 4–12). Each ACF was inverted according to the procedure suggested by Shusterman et al. (16, 17), to yield an apparent MSD, 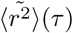, to be compared to the actual MSD. ACFs were also fitted to the general expression often used to assess anomalous diffusion processes, Eq. 15, in order to construct the diffusion law and recover an apparent anomalous exponent, *α*.

**Figure 4:**
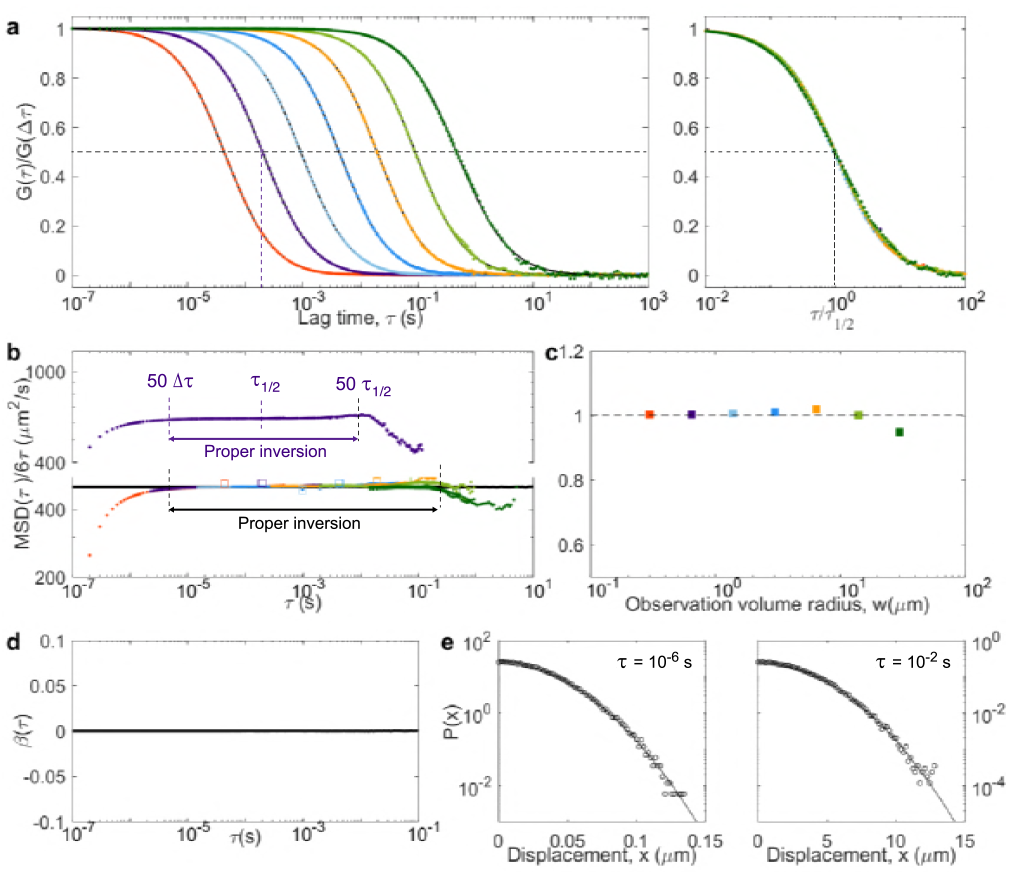
Simple diffusion (*D* = 500*µ*m/s^2^). (a) ACFs obtained for different detection volume sizes (coloured symbols: simulations; black lines: fit of Eq. 15 to the simulated data). The same ACF, with lag time normalized by *τ_1/2_*, <, are shown in the right panel. (b) Apparent diffusion coefficient, 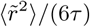, calculated from the inversion of the ACF (solid coloured symbols) and compared with the actual diffusion coefficient, (*r*^2^)/(6*τ*), obtained directly from the particle trajectories (black line). Empty symbols show the diffusion law (i.e. the value of *w*^2^/(4*τ*_1/2_)) as a function of τ_1/2_ for each ACF). One of the inverted curves is also shown on its own and shifted upwards. (c) Apparent anomalous exponent obtained from the fit of the ACFs in (a). The dashed line is a guide for the eye showing the expected *α* = 1 value for simple diffusion. (d) Non-gaussian parameter calculated from the trajectories (black line) and expected *β* = 0 value for simple diffusion (dashed line). (e) Distributions of displacements for *τ* = 10^−6^*s* (left) and *τ* = 10^−2^*s* (right). Lines are fit to a Gaussian distribution.

### 4.1 Simple Brownian diffusion

We first considered simple Brownian diffusion, to check how faithfully our simulations could reproduce analytical results (Fig. 4). As calculated from particle trajectories, the expected (*r*^2^) = *6D Τ* and *β* = 0 (black lines in Fig. 4b and d) are obtained with good precision up to τ = 10 s. The ACFs obtained from the simulated VLS-FCS data have the expected self-similar form, with a normal diffusion law (*w*^2^ *α τ*_1/2_, Fig. 4b) and *α* ~≃ 1 (Fig. 4c). Performing the ACF inversion in this simple Gaussian case, where we should have 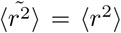 at all T, shows that three separate factors affect the quality of 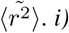 Due to the finite simulation time, for *τ* > 1 s the ACFs (and therefore 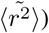 become noisy.*ii)* Because simulations were done for regular arrays of detection volumes spaced by 15 *w*, a small positive correlation ~ 0.1% of the ACF amplitude), resulting in a dip in 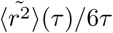, is observed around (15 *w*)^2^/4*D* = 225*τ_D_* (see section 2.6 and Fig. 2).*iii)* The apparent MSD 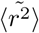 differs from (*r*^2^) at short lag times (*τ* < 50Δ*τ*) because of the imperfect normalization of the ACFs, which were just divided by their value at Δ*τ* = 10^−7^ s (see section 2.4 and Fig. 1). Despite these limitations, considering only the inverted ACFs between 50Δ*τ* and 10*τ_D_* for each ACF, results in a apparent MSD which is equal to the actual MSD for 5 decades in time, from below 10^−5^ s to beyond 10^−1^ s.

### 4.2 Continuous time random walk

We next simulated an anomalous diffusion model, a CTRW, where a wait time *τ_w_* drawn from a power law distribution 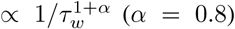 was introduced between each step. CTRW processes are non-ergodic (29, 35). As a consequence, the MSD obtained by performing both a time and ensemble average is linear in time (Fig. 5). However, when performing only an ensemble average, the MSD is anomalous, 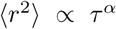 (Fig. 6). Our simulation shows that the non-Gaussian parameter also depends on how averages are performed: it is high (because of a large number of immobilized particles) and decays as a power law when performing a time average (Fig. 5d), but has a low and almost constant value when performing only an ensemble average (Fig. 6d). To check whether ergodicity can also be detected from FCS experiments, we calculated ACFs in two ways: first in the usual way (performing a time average on the signal, which already represents an ensemble average over the different particles present in the observation volume, Fig. 5), and second by averaging over many different repeats of the experiment instead of over time (ensemble average, Fig. 6, as also done in Ref. (35)). Whereas ACFs calculated in these two different ways are similar for ergodic processes, we see a clear difference for the CTRW process. The time-averaged ACFs never stabilize (their shape depends on the length of the measurement), a reflection of the ageing of the sample (36). Their shape also depends on the size of the observation volume (with *α* increasing from 0.5 to 1 as *w* increases). The apparent MSD extracted by inversion of these ACFs differs from the actual linear MSD and from one another (consistent with the fact that the process is strongly non-Gaussian), with 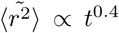 over a large range of lag times. Of note, however, apparent and actual MSD coincide when 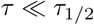 (a regime that can be observed only for the larger detection volumes), as expected from the calculations presented in section 2.3. In contrast, the ensemble-average ACFs (Fig. 6) are well-behaved and self-similar, with an apparent anomalous exponent *α* ≃ 0.7 close to the actual *α* = 0.8. Accordingly, the inversion of the ensemble-averaged ACFs gives apparent MSD that largely capture the power-law dependence of the actual MSD (except for those curves with very short T_1/2_ for which the normalization does not work well).

**Figure 5:**
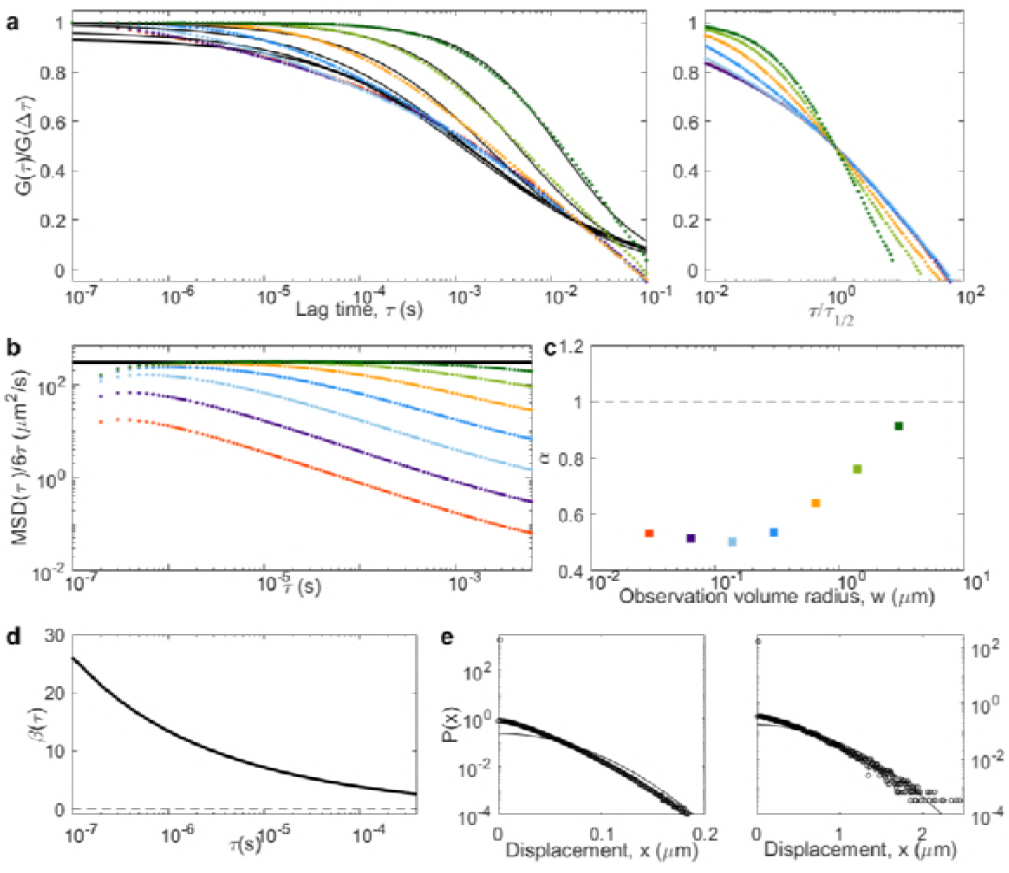
CTRW, where the ACFs (a), MSD (b), non-Gaussian parameter (d) and distribution of displacements (e) have been calculated in the same way as for all the other models, that is taking both a time and an ensemble average over time and particles. Panels are as in Fig. 4.

**Figure 6:**
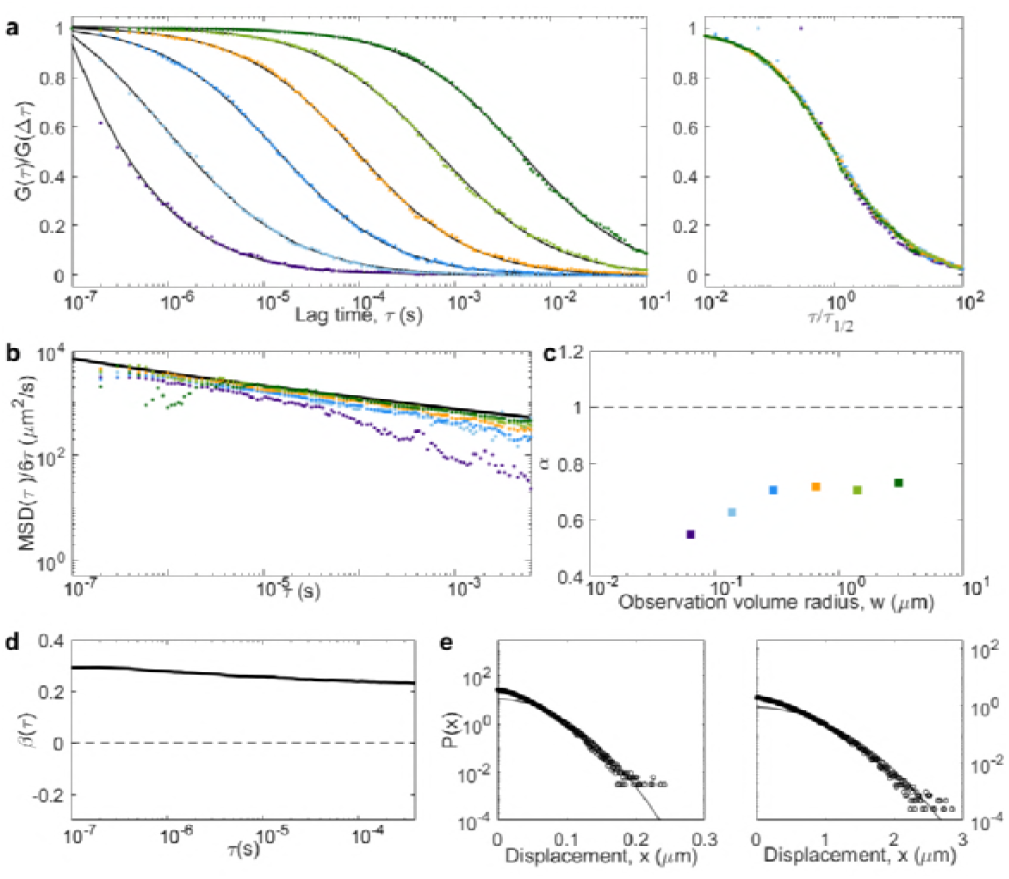
CTRW, where the ACFs (a), MSD (b), non-Gaussian parameter (d) and distribution of displacements (e) have been calculated using only an ensemble average over particles. Panels are as in Fig. 4.

Our simulations confirm the result from previous 2D simulations, which had shown that for a CTRW, the anomalous exponent recovered from FCS experiments using Eq. 15 can be significantly different than the real anomalous exponent (29, 37). In addition, our simulations emphasize that the apparent *α* depends strongly on the size of the observation volume (Fig. 5c), a phenomenon directly linked to the scale-dependence of the non-Gaussian parameter (Fig. 5d). Calculating the ACF as an ensemble-average only (as first done in Ref. (35)), although not necessarily easy to achieve experimentally, leads to much better behaved results, with stable self-similar ACFs, and an apparent anomalous exponent approaching the actual one.

### 4.3 Two-component diffusion

We next examined three cases in which tracer particles could switch (with constant rates *k*_on_ and *k*_off_) between two modes with diffusion coefficients *D* and *D’*.

The simplest case is that of two separate populations of tracers (fractions *f* and *f’* = 1 - *f*) that are not allowed to interchange (*k*_on_ = *k*_off_ = 0). Although a simple analytical solution exists in this case, we still performed a simulation, whose results are shown in Fig. 7. For such a process the MSD is linear in time, (*r*^2^} = 4 *D*_avg_τ at all lag times with *D*_avg_ = *fD* + (1 - *f*)*D’*. Yet the propagator associated with the process is non-Gaussian (it is the sum of two Gaussians). The non-Gaussian parameter has a constant value, 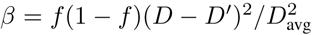. The ACFs (a weighted sum of the ACFs that would be obtained from either population of tracers) are self-similar, with an apparent *α* ~ 0.8 in the conditions of our simulation. Because of the non-Gaussian nature of the process, 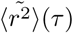 coincides with (*r*^2^}(*τ*) only for 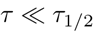. For 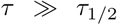, all the 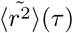 instead approach a simple diffusion MSD with diffusion coefficient 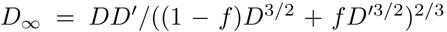. The diffusion law is linear in time, with an apparent diffusion coefficient comprised between *D*_avg_ and 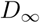.

**Figure 7:**
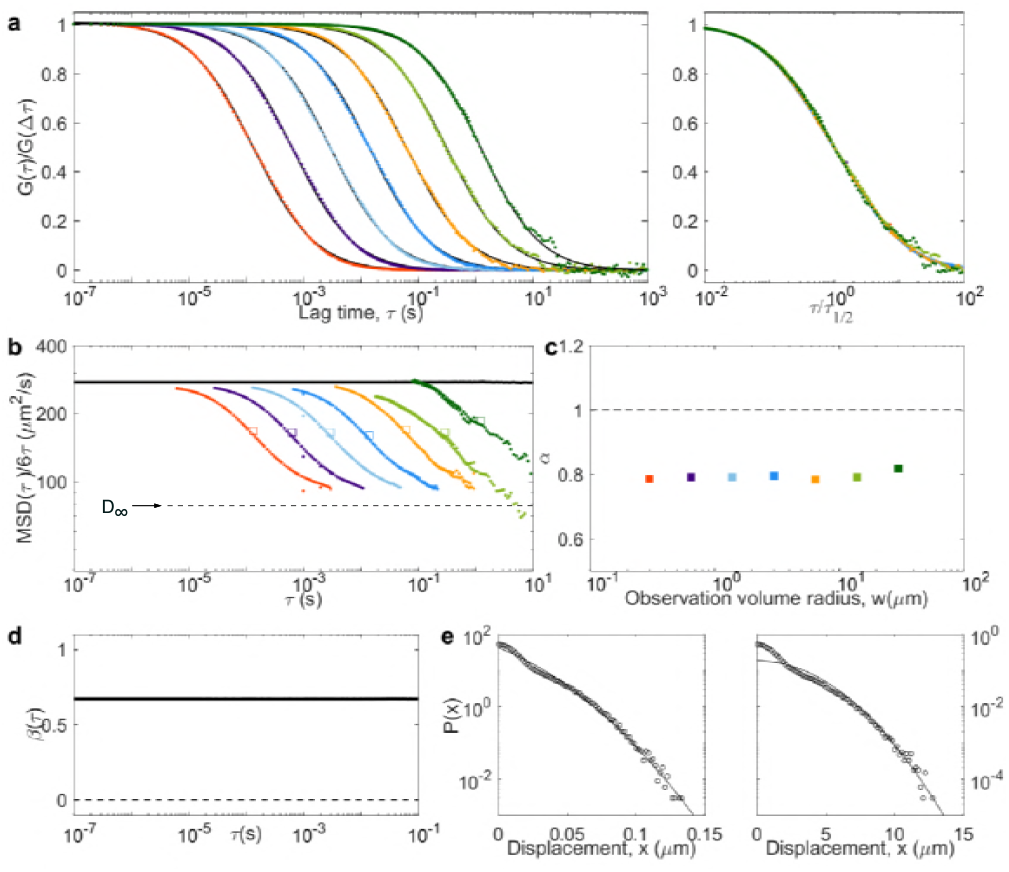
Two-component diffusion for separate populations of tracers (*k*_on_ = *k*_off_ = 0, *D* = 500*µm*^2^*/s, D’* = 50*µm^2^/s, f* = 0.5). In this case, one expects *D*_avg_ = 275*µ*m^2^/s, 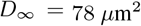 and *β* = 0.67.

We then considered the general case, where particles can switch back and forth between two simple diffusion modes with Poisson statistics (*k*_on_ = *k*_OFF_ = 500 *s*^−1^). In this scenario (one of the few considered before in the context of VLS-FCS (38, 39)), we expect a cross-over around the relaxation time, *τ*_C_ = 1 /(*k*_on_ + *k*_OFF_)). In the “fast diffusion” regime 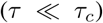 tracers remains in the same state while crossing the observation volume and behave as two separate populations. In the “fast reaction” regime 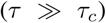 tracers switch state many times while crossing the observation volume, and appear to be undergoing simple diffusion with diffusion coefficient *D*_avg_. Meanwhile, *β* passes from 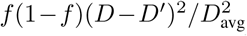 to 0 around *τ_c_*, and a becomes subdiffusive below 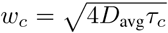 (Fig. 8). As in the previous case, the actual MSD is linear in time at all lag times. However, this time the propagator is Gaussian for *τ* > *τ_c_*, thus 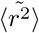 coincides with (*r*^2^} both below *τ*_1/2_ and above *τ_c_*, and tends towards D in between. Notably, for large w > w _c_ ((i.e. *τ*_1/2_ > *τ_c_*), 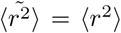 at all lag times.

**Figure 8:**
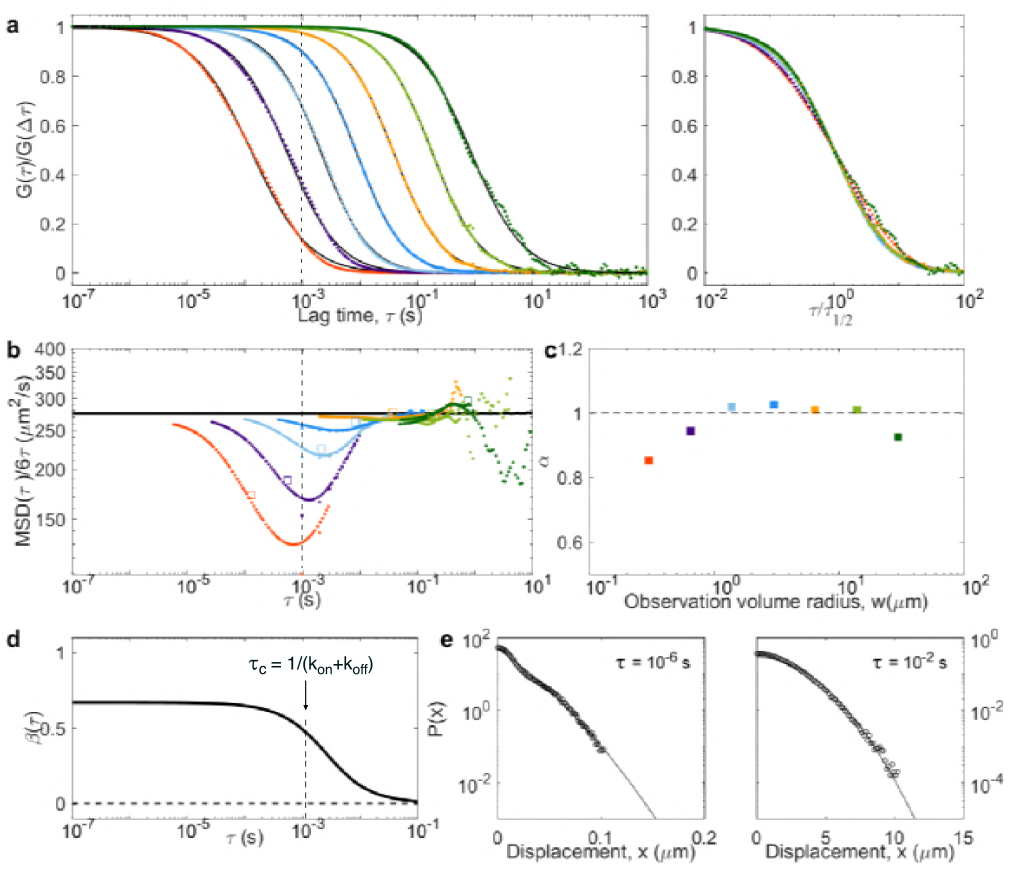
Two-component diffusion with *D* = 500*µ*m^2^/s, *D’* = 50*µ*m^2^/s and *k*_on_ = *k*_off_ = 500 *s*^−1^. In these conditions,a crossover between non-Gaussian and Gaussian diffusion is expected around *τ_c_* = 1ms and *w_c_* = 1 µm.

Finally, we considered the limiting case in which tracers transiently experience immobilization (*D’* = 0). The signature of this “stick-and-diffuse” model has been considered in the case of single-scale FCS experiments (40). It differs significantly from a CTRW because the distribution of immobilization times is exponential. Its VLS-FCS signature is no different from that of the general case, with a crossover from anomalous and non-Gaussian to Gaussian around the crossover time *τ*_C_ = 1 ms (Fig. 9), but with higher non-Gaussianity (*β* = (1 - *f)/f* at small T), lower *D*_avg_ = *fD* and 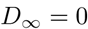.

**Figure 9:**
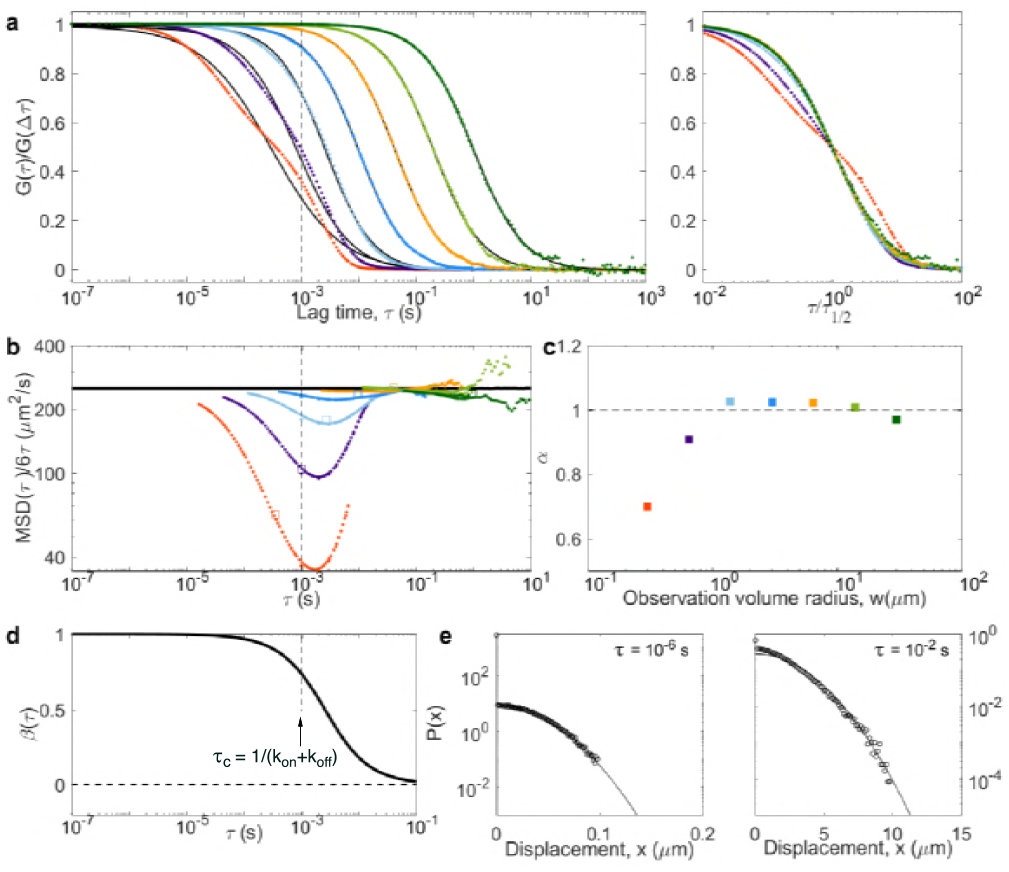
Two-component diffusion with *D* = 500*µ*m^2^/s and *D’* = 0*µ*m^2^/s (stick-and-diffuse). The transition rates were *k*_on_ = *k*_off_ = 500 *s*^−1^, thus *f* = 0.5. Then *D*_avg_ = 250*µ*m^2^/s, *D*= 0*µ*m^2^/s, and *β* = 1 at small*τ*. The characteristic time is τ_C_ = 1 ms, corresponding to *w*_c_ ~ 1*µ*m.

### 4.4 Diffusing diffusivity

As another example of a diffusive process with linear MSD but non-Gaussian propagator, we considered the diffusive diffusivity model (31). Particles were given a diffusion coefficient which varied in time according to a 1D random walk (with diffusion coefficient*d* = 5.5 x 10^7^*µ*m^4^/*s*^3^) between *D*_max_ = 500*µ*m^2^/*s* and *D*_min_ = 0*µ*m^2^/*s*. We then expect - and observe - a crossover around T_C_ = (*D*_max_ - *D*_min_)^2^/ (2 *d*) = 2 ms. The MSD is linear at all lag time, with apparent diffusion coefficient *D*_app_ = (*D*_max_ - *D*_MIN_))/2, and a switch from non-Gaussian (*β* ~ 0.35) to Gaussian (*β* = 0) behaviour around *τ_c_*(Fig. 10). Below *τ_c_*, the distribution of displacements approaches an exponential distribution, as previously noted by Chubynsky & Slater (31). Accordingly, the ACFs display an anomalous shape for w < *w_c_* ~ 1*µ*m, with *α* approaching 0.9 at the smallest detection volumes. As for the two-component models, 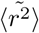 coincide with *(r^2^)* both for *τ* < *τ*_1/2_ and *τ* > *τ_C_*. Overall, the signature of diffusing diffusivity resembles that of a two-component model, but with less marked anomalous features.

**Figure 10:**
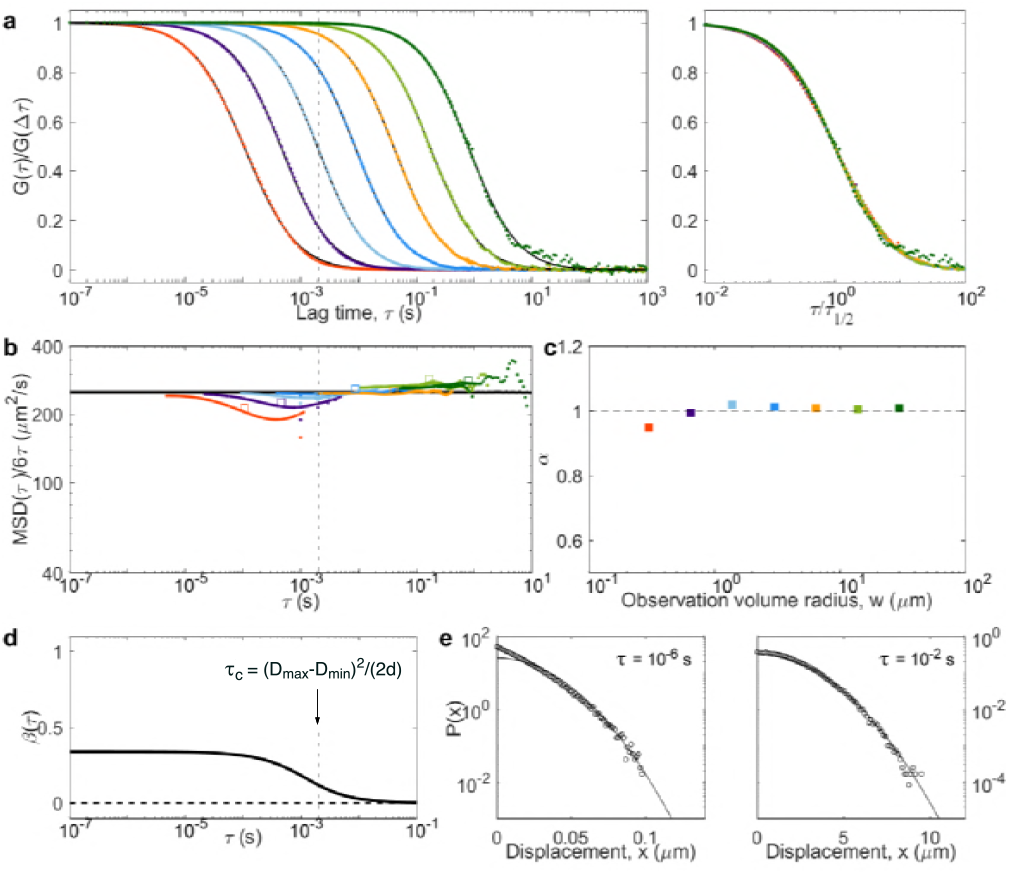
Diffusing diffusivity, with a diffusion coefficient diffusing between *D*_min_ = 0*µ*m^2^/s and *D*_max_ = 500*µ*m^2^/s with a diffusion coefficient*d* = 5.5 x 10^7^*µ*m^4^/*s*^3^.

### 4.5 Obstructed diffusion

Another model often invoked to account for anomalous diffusion in complex media is obstructed diffusion, where the motion of the tracers is restricted by the presence of immobile obstacles. If the obstacle concentration is below the percolation limit, anomalous diffusion occurs as a transient regime between unhindered short-scale diffusion and large-scale effective diffusion. We simulated obstructed diffusion using randomly placed cubic obstacles of size *L* at a volume fraction *ϕ* slightly below the percolation limit (*ϕ*\* = 0.3116 in this geometry). As seen before in 2D simulations (22, 41), the apparent diffusion coefficient of the tracers switches from their actual diffusion coefficient (*D* = 500*µ*m^2^/s) to an effective value (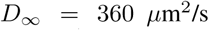 in our conditions) around *τ*_C_ = *L*^2^ /(6 *D*^2/3^) ~ 0.4 ms (Fig. 11). This transient anomalous regime is visible in the ACFs around 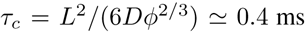, where has a minimum. Just above *τ_c_*, a small peak is observed for the non-Gaussianity factor (a similarly weak non-Gaussianity was shown for 2D simulation of obstructed diffusion at large lag time (42)). Accordingly, 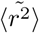 deviates very slightly from (*r*^2^) in that region, for curves for which *τ*_1/2_ is around or below τ _C_.

**Figure 11:**
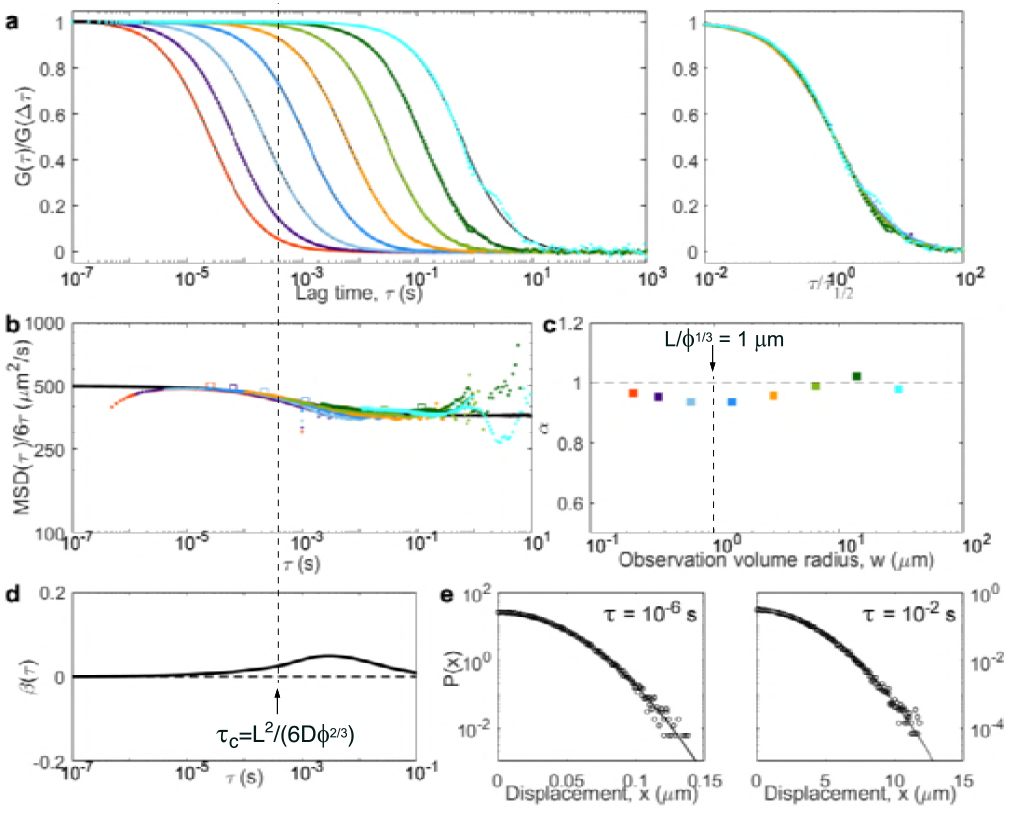
Obstructed diffusion, with *D* = 500*µ*m^2^/s, and cubic obstacles of size *L* = 0.75*µ*m at a volume fraction *ϕ* = 0.31.

### 4.6 Caged diffusion

The last considered model was caged diffusion, where tracer particles diffuse through an array of semi-permeable cages. As for obstructed diffusion, a transitory anomalous regime visible in the ACFs is expected (43–45). We simulated caged diffusion for regularly arranged cubic cages of size *L* = 1*µ*m, and a probability *p* = 0.005 to cross the barriers between cages when encountering them (Fig. 12). We observe the expected crossover between short-term and long-term diffusion coefficient, with a MSD that more or less follows the approximate 2D analytical expression derived by Destainville et al. (i.e. a switch from microscopic to effective diffusion coefficient around the relaxation time of the particles in the cages) (32, 44, 46). Interestingly, the non-Gaussian parameter displays two small amplitude peaks. The first, found below *τ_c_* = (*L*)^2^/6 *D* = 0.3 ms, corresponds to the particles equilibrating in a cage. The second corresponds to the particles leaving the cage. Yet as *β* remains small, 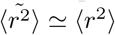 at all lag times (the strongest deviation is observed just above *τ_c_*). ACFs with *w* ~ *L* reflect the imperfect confinement of the particles in the cage by displaying two separate timescales, and therefore *α* < 1.

**Figure 12:**
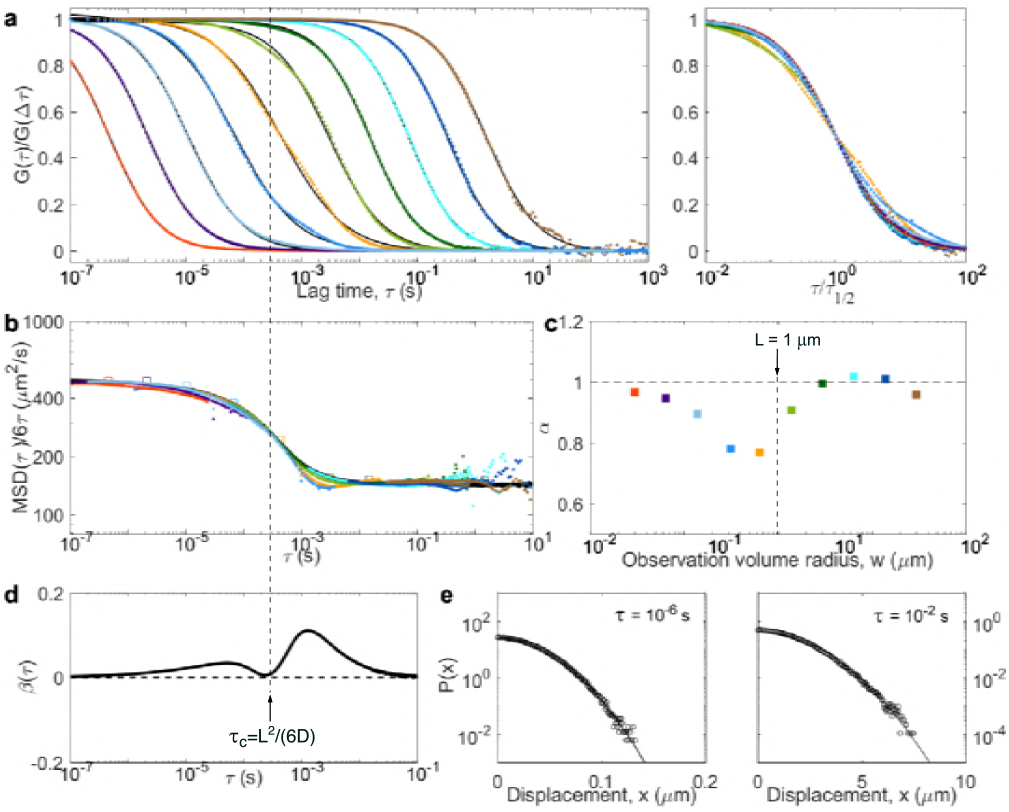
Caged diffusion (small escape probability), with *D* = 500*µ*m^2^/s, *L* = 1*µ*m and *p* = 0.005.

An interesting particular case is that of impermeable cages (*p* = 0). In practice, FCS experiments would be difficult to perform for such a system, as confined particles photobleach rapidly, yet it is interesting to think about the signature of such processes. In this case, the apparent diffusion coefficient falls all the way to 0 above *τ_c_*, and instead of a second peak, *β* goes to negative values (data not shown). The distribution of displacements has a hard limit at *x* = *L*, eventually assuming a triangular shape at large *τ*. As w increases, the ACFs rapidly assume a shape reflecting confinement, with *α* > 1 and with a characteristic decay time *τ*_1/2_ that no longer depends on *w*.

## 5 Discussion

Each of the models considered in this work is representative of a class of diffusion processes relevant to the cellular environment. CTRW and two-component diffusion are models of diffusing molecules interacting with slow or immobile binding partners, while diffusing diffusivity, obstructed diffusion and caged diffusion reflect different crowding scenarios. In cells, crowding and molecular interactions both play a part in protein mobility. Our goal was thus to determine how well the influence of these different scenarios could be distinguished in VLS-FCS experiments.

Until recently, the information contained in VLS-FCS experiments has been exploited via the dependence of τ_1/2_ on *w*^2^ (diffusion law), initially introduced to distinguish simple diffusive processes (*τ*_1/2_ = = w^2^) from photophysical processes (*τ*_1/2_ independent of *w*) (47). This model-independent approach has also proved useful to help distinguishing between different types of diffusion (5, 7, 13, 48). For example, in 2D, caged diffusion and dynamic partitioning into domains result in negative and positive τ = 0 intercepts of the diffusion law, respectively (13, 43). However, our 3D VLS-FCS simulations illustrate an important limitation of the diffusion law, which is that it coincides with the MSD only for Gaussian or near-Gaussian processes. In more complex cases (CTRW, two-component diffusion), the diffusion law relates to the MSD in a non-trivial and ill-defined way, thus one must make assumptions about the underlying diffusion process to extract information from it (a problem already pointed out in the context of imaging FCS (49)).

Ultimately, the issue with the diffusion law is that it collapses the rich information contained in the shape of the ACF into a single value, *τ*_1/2_. In contrast, analysis of the detailed shape of the ACFs obtained at different *w* can give a lot of information about the underlying process (as shown for obstructed diffusion (22), two-component diffusion (38, 39) or diffusion in phase separated membranes (50)). However, it is model-dependent. This is why the inversion procedure introduced by Krichevsky et al. to obtain the MSD from the ACF (see section 3.5) (16, 17), and its combination with lengthscale variation as first considered by Hofling and Fra-nosch (22), and later experimentally demonstrated by us (20), is so powerful. It uses the full range of information contained in the ACFs to allow a precise characterization of the diffusion process, over many decade in times, in a model-independent manner.

By using the same simulation framework to study different classes of diffusion models and by simulating VLS-FCS experiments, we provide here a library of inverted ACFs, and thus an unbiased way to interpret the results of such experiments. Although comparison of different anomalous diffusion models using simulations are available (notably a comparison of fractional Brownian motion and CTRW (29)), none has ever been examined from the point of view of inverted VLS-FCS. All the considered models (except for simple diffusion) resulted in ACFs with “anomalous” behaviour, i.e. with *α* < 1 for at least some of the observation volume sizes considered. However, they varied greatly with respect to several essential features that can be accessed through a VLS-FCS experiment, namely: 1) self-similarity vs. presence of a characteristic timescale (visible as a crossover in the inverted ACFs), 2) linearity of the MSD, 3) Gaussianity of the distribution of displacement. All these models are non-Gaussian, an important characteristic observed in many biological systems (51), e.g. RNA-protein particle diffusion in cells (52). Of particular interest, the models involving interchange between different diffusion modes with Poisson statistics (two-component models) or via a continuous diffusive process (diffusing diffusivity) exhibit an “anomalous, yet Brownian” behaviour (non-Gaussian propagators associated with a “normal” linear MSD), a feature observed in many biologically relevant contexts, for example diffusion in crowded polymer solutions (20) or actin networks (53). This linear behaviour of the MSD can be observed in inverted VLS-FCS, whereas the diffusion law has an apparent power-law dependence on time (Figs. 8–10).

It has been pointed out that a limitation of the inversion procedure (the fact that it faithfully returns the MSD only for Gaussian processes), could be turned to an advantage in the case of VLS-FCS, as it can be used as a test for Gaussianity (20, 23). Indeed, for all the models considered here, at those lag times for which the process is not Gaussian (i.e. where 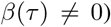, we observe a spread in the value of the inverted ACFs obtained for different w (i.e. 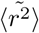 depends on w). Moreover, the further *β* (*τ*) is from 0, the further the family of inverted ACFs deviate from one another. More than a simple test for Gaussianity, an inverted VLS-FCS can thus inform on the range of lag times over which the process is non-Gaussian, and on how far from Gaussian the process is.

Maybe the most interesting result from our study is that, regardless of Gaussianity, the inversion procedure based on Eq. 3 returns the actual MSD if 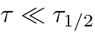. This can be proven by performing a Taylor expansion of the ACF in (*r*^2^} /*w*^2^ (as shown in section 2.3), and can be seen in for all the models simulated here. The linear relationship existing between the ACF and the MSD at short lag time has been noted before (24). However, we show here that this linear regime can be greatly extended by increasing w. Performing a single-point FCS experiment for a large detection volume, will allow recovering the actual MSD from « 20Δ*τ* (where Δ*τ* is the experiment’s time resolution) up to ~ *τ*_1/2_/10, whether the underlying process is Gaussian or not. Of course, this will hold true only if good statistics can be achieved (i.e. the studied system is stable enough to allow long measurements) and if the photophysics of the dye used to tag the tracer is not an issue at short *τ*. The MSD can then be retrieved for all kinds of diffusive processes, over several orders of magnitude in time (something which we have previously observed for fluorescent beads diffusing in gels (20)). Remarkably, the MSD can then extend below the diffraction limit (a point which has been made before eloquently (14, 54)). Thus, in FCS experiments, the best strategy to retrieve information on processes at short time scale may not be to physically achieve subdiffraction-limited detection volume, but instead to work on achieving high quality PSF for larger detection volumes with a shape as close as possible to a 3D Gaussian profile (ensuring the inversion procedure faithfully returns the MSD at short lag time).

## 6 Author Contributions

MS designed and performed simulations, helped write the manuscript and prepared figures. CF helped design simulations and wrote the manuscript.

## 7 Acknowledgments

This article is dedicated to Jörg Langowski, whose encouragements have been an inspiration for our work, and who was very passionate about the capacity of FCS to resolve subdiffraction motions. We also thank Felix Hofling for many helpful discussions. This work was funded by the Natural Sciences and Engineering Research Council of Canada (NSERC), and enabled by the use of computing resources provided by SHARCNET and Compute Canada.

